## Supplementary material for "Gene body DNA hydroxymethylation restricts the magnitude of transcriptional changes during aging": Occean_combined_supp_v2

**Supp. Figure 1: Quality control metrics of mouse liver hMeDIP-seq libraries**

(**A**) (Left) Representative Bio-analyzer electropherogram of liver hMeDIP-seq library from young input and 5hmC samples. (Right), same as (left) but for old liver samples. (**B**) qPCR analysis for exogenous unmethylated and hydroxymethylated DNA control spike-ins. (**C**) Sequencing depth of 5hmC and input libraries from young and old liver samples. (**D**) Percent of alignment rates of 5hmC and input libraries to the GRCm38/mm10 genome. (**E**) The number of fragments mapped to the GRCm38/mm10 genome. (**F**) The length of mapped fragments extracted from SAM file. (**G**) LISA MOTIF analysis using the top 500 genes in young and old (ranked by gene body 5hmC signal). The top 20 young and old TFs (sorted by *p*-value (old x young)) are labeled. (**H**) Spearman-rank correlation between young (panel 1) and old (panel 2) gene body 5hmC signal and mRNA fold change (old vs young). Correlation between old vs young mean gene body 5hmC signal and mRNA fold change (panel 3). Correlation between random gene body 5hmC values and mRNA fold change (old vs young). ρ = spearman correlation coefficient.

**Supp. Figure 2: Quality control metrics of mouse liver MeDIP-seq libraries**

(**A**) (Left) Representative Bio-analyzer electropherogram of liver MeDIP-seq library from young input and 5mC samples. (Right) same as (left) but for old liver samples. (**B**) qPCR analysis for exogenous unmethylated and hydroxymethylated DNA control spike-ins. (**C**) Sequencing depth of 5mC and input libraries from young and old liver samples. (**D**) Percent of alignment rates of 5mC and input libraries to the GRCm38/mm10 genome. (**E**) The number of fragments mapped to the GRCm38/mm10 genome. (**F**) The length of mapped fragments extracted from SAM file.

**Supp. Figure 3: Age-related differences in 5hmC occur without detectable differences in 5mC**

(**A**) PCA plot obtained using input subtracted 5mC bigWig files. (**B**) Number of 5mC peaks called for young and old liver samples by MACS2 at q = 0.001. Data are summarized as mean ± standard deviation. (**C**) Violin plots (with boxplots) of peak characteristics size in bp (left), sum (middle), and mean (right) of signal. For $\log_2$(size) mean, young = 9.01, old = 9.02; $\log_2$(sum), young = 12.45, old = 12.53; $\log_2$(mean), young = 3.44, old = 3.50. *** = $p < 0.001$ by Welch's t-test. (**D**) Correlation heatmap of 5mC peaks called by MACS2 produced using DiffBind. (**E**) Volcano plot of differentially methylated regions (DMRs) in the liver identified by

DiffBind using $p < 0.05$. Hypo DMRs (fold change < 0) are peaks with less enrichment in the old and hyper DMRs (fold change > 0) are peaks with higher enrichment in the old. (**F**) Metaplots of 5mC signal at the DMRs in (E). (**G**) Metaplots of 5mC signal at the liver DHMRs in Fig. 2E. (**H**) Example genome browser tracks for liver hyper DMRs (intergenic and *Aimp1*) and hypo DMRs (*Fgfr1op2* and *Samd4*). (**I**) GO terms associated with the DMRs from (E) using GREAT. The top 5 biological process terms with FDR < 0.05 are shown. (**J**) Pie charts showing CpG and genic/intergenic annotations of the DMRs from (E). (**K**) Metaplot of 5mC signal over the gene bodies of all mm10 genes with quantifications for each replicate on the side. N/S = $p > 0.05$ by Welch's t-test.

**Supp. Figure 4: Supplementary for alternative splicing mediates 5hmC's transcriptionally restrictive function through decreased binding of splicing factors**

(**A**) Genome browser views of young and old 5hmC and 5mC signal at endogenous regions used to design oligo 1 (left) and oligo 2 (right). (**B**) Overlap between oligo 1 and oligo 2 for 5hmC-enriched proteins (top panel) and input-enriched proteins (bottom panel) when comparing 5hmC vs input. (**C**) GO analysis using DAVID for proteins overlapping between oligo 1 and oligo 2 in (B). (**D**) Heatmap (left) and volcano plot (right) of differential proteins ($p < 0.05$) in oligo 1 for the comparison of input-subtracted old 5hmC vs old 5mC and C. (**E**) Heatmap (left) and volcano plot (right) of differential proteins ($p < 0.05$) in oligo 1 for the comparison of input-subtracted old 5hmC vs young 5mC and C. (**F**) Heatmap (left) and volcano plot (right) of differential proteins ($p < 0.05$) in oligo 2 for the comparison of input-subtracted old 5hmC vs old 5mC and C. (**G**) Heatmap (left) and volcano plot (right) of differential proteins ($p < 0.05$) in oligo 2 for the comparison of input-subtracted old 5hmC vs young 5mC and C. (**H**) PCA plot of long-read sequencing data based on normalized transcript counts. (**I**) Examples of alternative splicing events in genes with high 5hmC. Shaded area shows a differentially used exon.

**Supp. Figure 5: Supplementary for quiescence and senescence drive the increase of 5hmC with age and impact cellular function**

(**A**) Dot blot for 5hmC signal in serum starvation-induced quiescent HepG2 cells. + control is 25 ng of young mouse hippocampus gDNA and – control is water. Quantifications are depicted below. **(B)** Senescence-associated β-galactosidase staining of proliferating cell and etoposide-, irradiation-, and oxidative stress-induced senescent cells. (**C**) qPCR analysis of *p16*, *p21*, *Lamin B1*, and *IL6*. ** = $p$ <0.01 and *** = $p$ < 0.001 by multiple t-test with FDR correction (two-stage step-up (Benjamini, Krieger, and Yekutieli). (**D**) BrdU assay of proliferating cells and etoposide-, irradiation-, and oxidative stress-induced senescent cells. Data are summarized as mean ± standard deviation. *** = q < 0.001 by multiple t-test with FDR correction (two-stage step-up (Benjamini, Krieger, and Yekutieli). (**E**) DHE staining of young and old mouse liver sections. 5hmC signal quantifications for mean intensity per nucleus is shown on the right. *** = $p$ < 0.001 by Welch's t-test. (**G**) DHE staining of HepG2 cells treated with $H_2O_2$ at indicated concentrations for 2 h. (**H**) DHE staining of HepG2 cells treated with 600 μM $H_2O_2$ for 2 h without NAC, sequential 24 h treatment with NAC after 2 h $H_2O_2$ treatment, or 2 h co-treatment with NAC followed by 24 h treatment with NAC only. (**I**) 5hmC dot blot of HepG2 cells from (H). Quantifications are depicted below. (**J**) 5hmC dot blot of HepG2 cells treated with $H_2O_2$ for 24 h. Quantifications are depicted below. For (A) and (I-J), two-way ANOVA with Tukey's multiple comparisons test yielded no significant results.

**Supp. Figure 6: Quality control metrics of mouse cerebellum hMeDIP-seq libraries**

(**A**) (Left) Representative Bio-analyzer electropherogram of cerebellum hMeDIP-seq library from young input and 5hmC samples. (Right) same as (left) but for old cerebellum samples. (**B**) qPCR analysis for endogenous controls, *Gapdh* (negative control) and *Sfi1* (positive control). (**C**) Sequencing depth of 5hmC and input libraries from young and old cerebellum samples. (**D**) Percent of alignment rates of 5hmC and input libraries to the GRCm38/mm10 genome. (**E**) The number of fragments mapped to the GRCm38/mm10 genome. (**F**) The length of mapped fragments extracted from SAM file.

**Supp. Figure 7: 5hmC's transcriptionally restrictive function extends to mouse cerebellum**

(**A**) PCA plot obtained using input subtracted 5hmC bigWig files. (**B**) Number of 5hmC peaks called for young and old cerebellum samples by MACS2 at q = 0.001. Data are summarized as mean ± standard deviation. (**C**) Violin plots (with boxplots) of peak characteristics size in bp (left), sum (middle), and mean (right) of signal. For $\log_2$(size) mean, young = 9.31, old = 9.36; $\log_2$(sum), young = 12.84, old = 12.93; $\log_2$(mean), young = 3.535, old = 3.571. *** = $p$ < 0.001 by Welch's t-test. (**D**) Correlation heatmap of 5hmC peaks called by MACS2 produced using DiffBind. (**E**) Volcano plot of DHMRs in the cerebellum identified by DiffBind using $p < 0.05$. Hypo DHMRs (fold change < 0) are peaks with less enrichment in the old and hyper DHMRs (fold change > 0) are peaks with higher enrichment in the old. (**F**) Metaplots of 5hmC signal at the DHMRs in (E). (**G**) Example genome browser tracks for cerebellum hyper DHMRs (*Pcdhga1* and *Plcb4*) and hypo DHMRs (*Rtn4r* and *Neurl1a*). (**H**) GO terms associated with the DHMRs from (E) using GREAT. The top 5 biological process terms with FDR < 0.05 are shown. (**I**) Pie charts showing CpG and genic/intergenic annotations of the DHMRs from (E). (**J)** Metaplot of 5hmC signal over the gene bodies of all mm10 genes with quantifications for each replicate on the side. N/S = $p > 0.05$ by Welch's t-test.

**Supp. Figure 8: Gene body 5hmC restricts the magnitude of transcriptional changes during cerebellum aging**

(**A**) Metaplots of 5hmC signal over the gene bodies of genes with low (n = 5900), intermediate (n = 5900), and high (n= 5900) average mRNA counts for young (left) and old (right) samples. (**B**) Metaplots of cerebellum 5hmC signal over gene bodies of genes with low (left, n = 5900) and high (right, n = 5900) |mRNA fold change| with age. Quantifications are depicted below. * = p < 0.05 by Welch's t-test. (**C**) Boxplots showing distribution of 5'UTR, 3'UTR, transcript, and CDS length as well number of exons for the for the genes with low (left) and high (right) |mRNA fold change| with age. *** = $p < 0.001$ by Welch's t-test. (**D**) Heatmaps of two GO terms from Supp. Fig. 2G showing gene body 5hmC signal and corresponding |mRNA fold change (old vs young)|. Heatmap is sorted by decrease in magnitude of mRNA fold change between old and young.

**Supp. Figure 9: Supplementary for human tissues also show 5hmC-mediated transcriptional restriction**

**(A)** Distribution of sex and age groups among GTEx donors used in this study. (**B**) (left) Heatmap output from ImpulseDE2 with monotonous and transiently changing genes with age in the brain. (middle) normalized mRNA count of brain-specific genes and brain-differential genes from (left). (right) GO plots associated with the brain-specific genes and brain-differential genes. (**C**) (left) Heatmap output from ImpulseDE2 with monotonous and transiently changing genes with age in the heart. (middle) normalized mRNA count of heart-specific genes and heart-differential genes from (left). (right) GO plots associated with the heart-specific genes and heart-differential genes. (**D**) (left) Heatmap output from ImpulseDE2 with monotonous and transiently changing genes with age in the liver. (middle) normalized mRNA count of liver-specific genes and liver-differential genes from (left). (right) GO plots associated with the liver-specific genes and liver-differential genes. Data are summarized as mean ± standard error of mean. ** = $p < 0.01$ and *** = $p < 0.001$ signifies whether mRNA counts are differential across any age groups by ANOVA test (without post-hoc comparisons).

**Supp. Figure 10: 5hmC is downregulated in response to high-fat diet and disulfiram**

(**A**). Schematic of diet regimen for standard and high fat diet. (**B**) Dot blot for 5hmC signal in gDNA isolated from mice liver fed on either a standard diet or high fat diet. + control is 200 ng of young mouse hippocampus gDNA and – control is water (**C**) 5hmC quantification from dot blot (B) stratified by type of diet. *** = $p < 0.001$ by two-way ANOVA with Šídák's multiple comparisons test. (**D**) Same as (C) except stratified by type of diet and sex. (**E**) Schematic of diet regimen for high fat diet and disulfiram drug treatment. (**F**) Dot blot for 5hmC signal in gDNA isolated from mice liver according to the procedure outlined in (E). Note that the bottom 3 samples for each column are females. (**G**) (Left) Quantification of 5hmC signal from (F) stratified by type of diet group. (Right) same as (left) except stratified by type of diet group and sex. Data are summarized as mean ± standard deviation. ** = $p < 0.01$, *** = $p < 0.001$ by two-way ANOVA with Tukey's multiple comparisons test.

**Supplementary Table Titles**

**Supp. Table 1: Animal information**

Sheet 1: DNA mass spectrometry

Sheet 2: Dot blot

Sheet 3: Immunofluorescence microscopy

Sheet 4: Liver hMeDIP and MeDIP

Sheet 5: RNA-seq reported elsewhere (Yang et al., *Molecular Cell*, under revision)

Sheet 6: Oligonucleotide mass spectrometry

Sheet 7: Dihydroethidium fluorescence

Sheet 8: direct RNA-seq

Sheet 9: Cerebellum hMeDIP and RNA-seq

Sheet 10: High fat diet mice obtained from Jackson Labs

Sheet 11: High fat diet and disulfiram mice from Bernier et al. 2020

**Supp. Table 2: DNA mass spectrometry quantification from young and old tissues**

**Supp. Table 3: Liver differentially hydroxymethylated and methylated regions**

Sheet 1: Hypo DHMRs (old vs young)

Sheet 2: Hyper DHMRs (old vs young)

Sheet 3: Hypo DMRs (old vs young)

Sheet 4: Hyper DMRs (old vs young)

**Supp. Table 4: Liver RNA-seq gene rankings**

Sheet 1: Young - low mRNA group (DESeq2 normalized counts)

Sheet 2: Young – intermediate mRNA group (DESeq2 normalized counts)

Sheet 3: Young – high mRNA group (DESeq2 normalized counts)

Sheet 4: Old - low mRNA group (DESeq2 normalized counts)

Sheet 5: Old – intermediate mRNA group (DESeq2 normalized counts)

Sheet 6: Old – high mRNA group (DESeq2 normalized counts)

Sheet 7: low mRNA fold change (old vs young)

Sheet 8: high mRNA fold change (old vs young)

Sheet 9: low mRNA absolute fold change (old vs young)

Sheet 10: high mRNA absolute fold change (old vs young)

**Supp. Table 5: Oligo mass spectrometry protein quantification**

Sheet 1: Raw data

Sheet 2: Processed data

Sheet 3: Simplified data

**Supp. Table 6: Oligo mass spectrometry differentially enriched proteins**

Sheet 1: Oligo 1 - differentially enriched proteins (old 5hmC vs old input)

Sheet 2: Oligo 2 - differentially enriched proteins (old 5hmC vs old input)

Sheet 3: Oligo 1 - differentially enriched proteins (old 5hmC vs old 5mC and C)

Sheet 4: Oligo 1 - differentially enriched proteins (old 5hmC vs young 5mC and C)

Sheet 5: Oligo 2 - differentially enriched proteins (old 5hmC vs old 5mC and C)

Sheet 6: Oligo 2 - differentially enriched proteins (old 5hmC vs young 5mC and C)

**Supp. Table 7: Alternative splicing results from rMATS**

Sheet 1: Skipped exon alternative events – inclusion difference (old vs young)

Sheet 2: Alternative 5’ splice site events – inclusion difference (old vs young)

Sheet 3: Alternative 3’ splice site events – inclusion difference (old vs young)

Sheet 4: Mutually exclusive exon events – inclusion difference (old vs young)

Sheet 5: Retained intron events – inclusion difference (old vs young)

**Supp Table 8: Differential isoform usage from direct RNA-seq**

**Supp Table 9: Transcript length data using Nanoplen**

**Supp. Table 10: Cerebellum differentially hydroxymethylated regions and RNA-seq gene rankings**

Sheet 1: Hypo DHMRs (old vs young)

Sheet 2: Hyper DHMRs (old vs young)

Sheet 3: Young - low mRNA group (DESeq2 normalized counts)

Sheet 4: Young – intermediate mRNA group (DESeq2 normalized counts)

Sheet 5: Young – high mRNA group (DESeq2 normalized counts)

Sheet 6: Old - low mRNA group (DESeq2 normalized counts)

Sheet 7: Old – intermediate mRNA group (DESeq2 normalized counts)

Sheet 8: Old – high mRNA group (DESeq2 normalized counts)

Sheet 9: low mRNA absolute fold change (old vs young)

Sheet 10: high mRNA absolute fold change (old vs young)

**Supp. Table 11: GTEx and ImpulseDE2 results**

Sheet 1: List of age-related differential genes in brain identified from ImpulseDE2

Sheet 2: List of age-related differential genes in heart identified from ImpulseDE2

Sheet 3: List of age-related differential genes in liver identified from ImpulseDE2

Sheet 4: List of brain-differential and brain-specific genes

Sheet 5: List of heart-differential and heart-specific genes

Sheet 6: List of liver-differential and liver-specific genes

**Supp. Table 12: Antibodies and DNA oligo sequence**

Sheet 1: Antibodies

Sheet 2: DNA oligo sequence used for oligo mass spectrometry

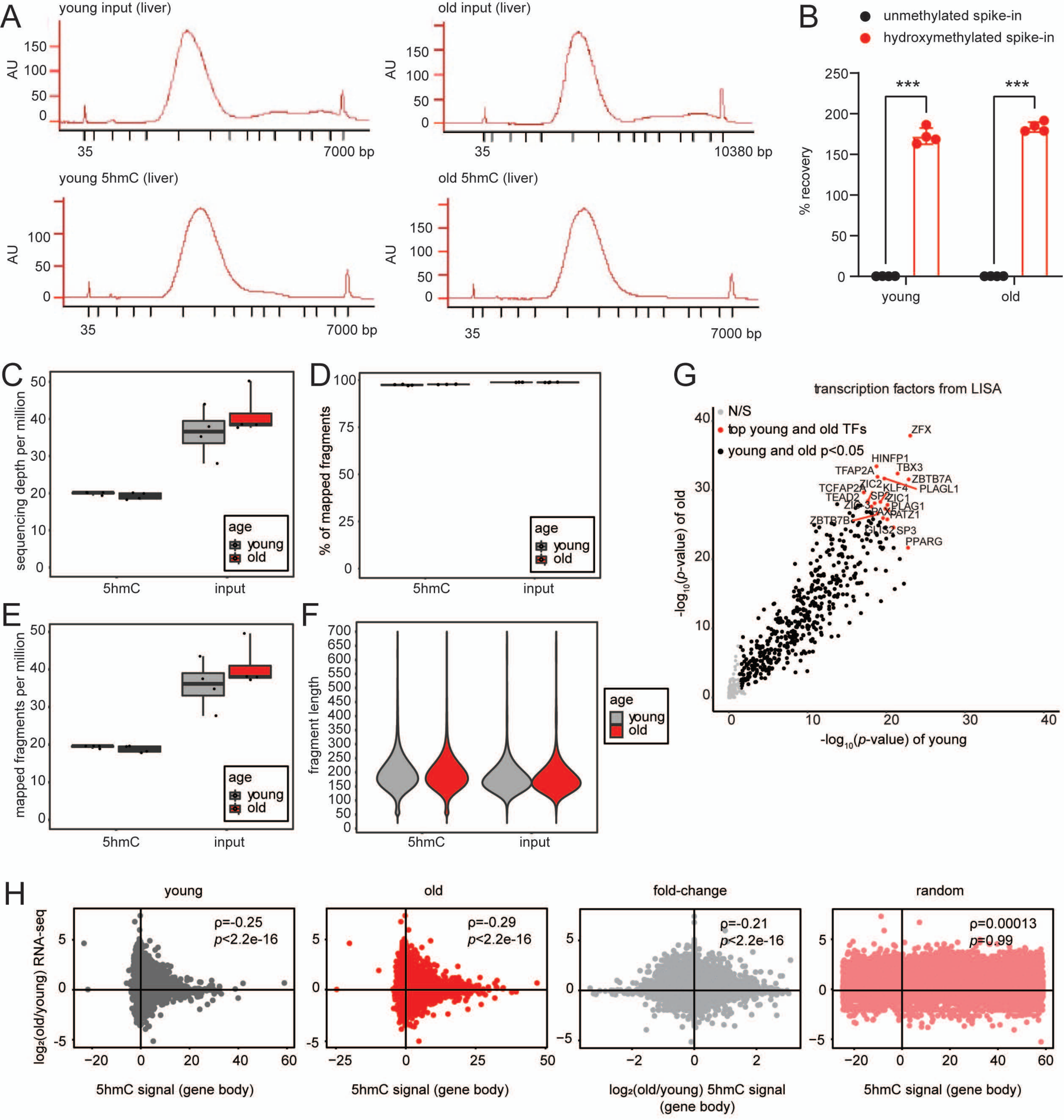

A
young input (liver)
old input (liver)
young 5hmC (liver)
old 5hmC (liver)
AU
35
7000 bp
10380 bp
B
unmethylated spike-in
hydroxymethylated spike-in
% recovery
***
young
old
C
sequencing depth per million
age
young
old
5hmC
input
D
% of mapped fragments
E
mapped fragments per million
F
fragment length
G
transcription factors from LISA
N/S
top young and old TFs
young and old p<0.05
ZFX
HINFP1
TBX3
TFAP2A
ZBTB7A
PLAGL1
TCFAP2A
ZIC2
KLF4
TEAD2
SP2
ZIC1
PLAG1
ZIC3
PAX5
PATZ1
ZBTB7B
GLIS2
SP3
PPARG
-log10(p-value) of old
-log10(p-value) of young
H
young
old
fold-change
random
ρ=-0.25
p<2.2e-16
ρ=-0.29
p<2.2e-16
ρ=-0.21
p<2.2e-16
ρ=0.00013
p=0.99
log2(old/young) RNA-seq
5hmC signal (gene body)
log2(old/young) 5hmC signal (gene body)


Figure S1

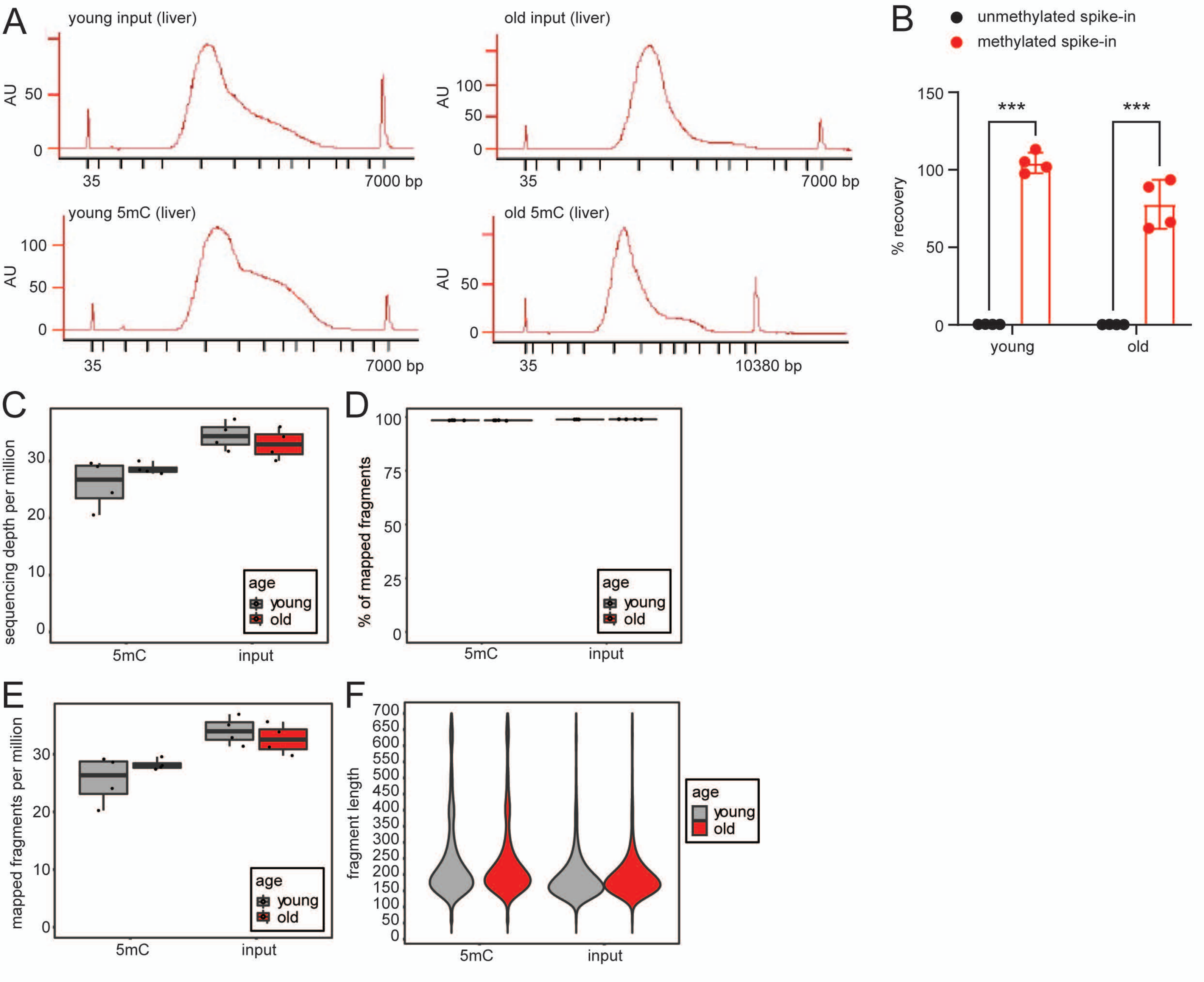
A
young input (liver)
old input (liver)
young 5mC (liver)
old 5mC (liver)
AU
35
7000 bp
10380 bp
B
unmethylated spike-in
methylated spike-in
% recovery
***
young
old
C
sequencing depth per million
age
young
old
5mC
input
D
% of mapped fragments
E
mapped fragments per million
F
fragment length

Figure S2

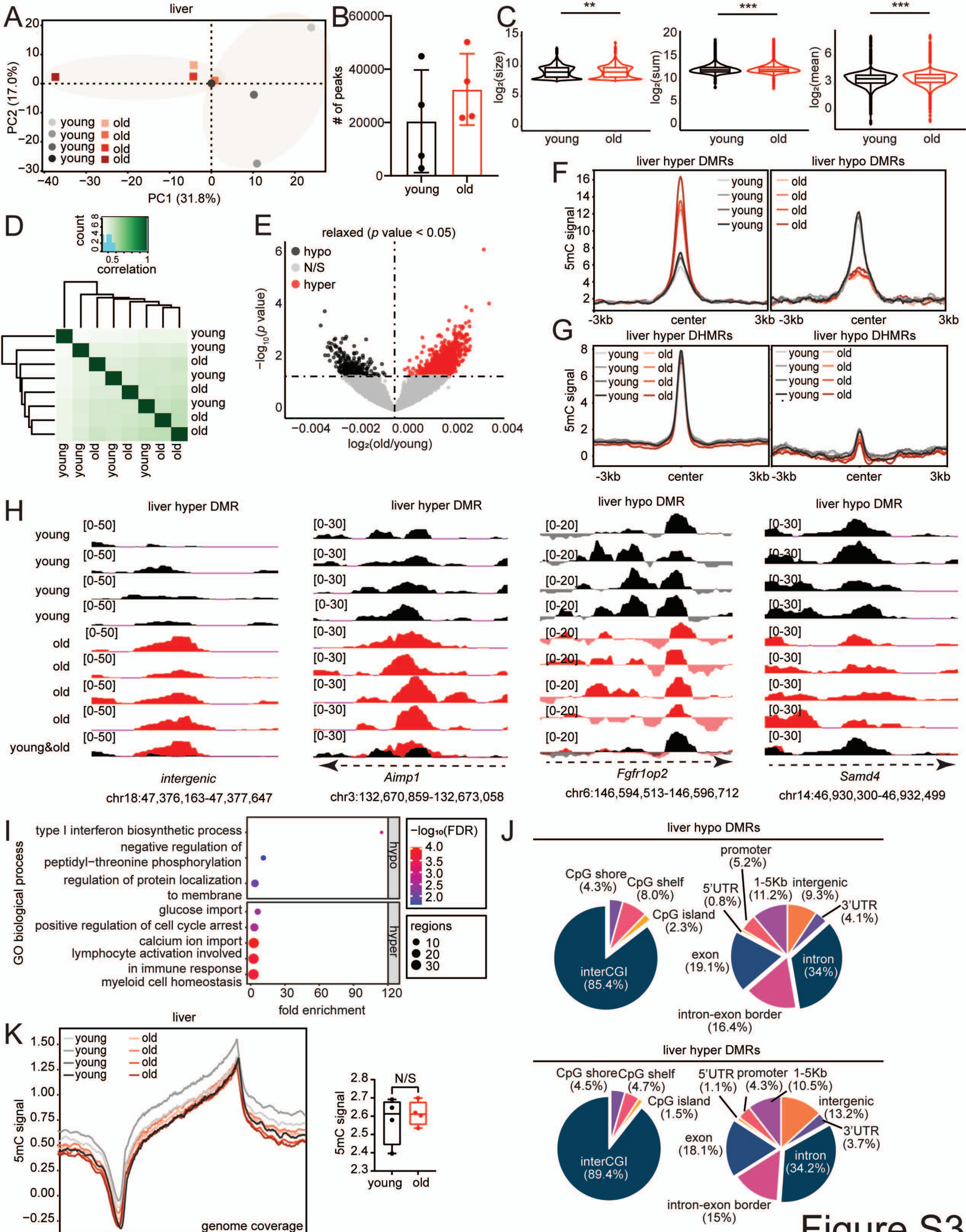
A
liver
PC2 (17.0%)
PC1 (31.8%)
young
old
B
### of peaks
young
old
C
log2(size)
log2(sum)
log2(mean)
young
old
**
***
***
D
count
correlation
young
old
E
relaxed (p value < 0.05)
hypo
N/S
hyper
−log10(p value)
log2(old/young)
F
liver hyper DMRs
liver hypo DMRs
5mC signal
-3kb
center
3kb
G
liver hyper DHMRs
liver hypo DHMRs
5mC signal
H
liver hyper DMR
liver hypo DMR
young&old
intergenic
chr18:47,376,163-47,377,647
Aimp1
chr3:132,670,859-132,673,058
Fgfr1op2
chr6:146,594,513-146,596,712
Samd4
chr14:46,930,300-46,932,499
I
GO biological process
type I interferon biosynthetic process
negative regulation of peptidyl−threonine phosphorylation
regulation of protein localization to membrane
glucose import
positive regulation of cell cycle arrest
calcium ion import
lymphocyte activation involved in immune response
myeloid cell homeostasis
fold enrichment
−log10(FDR)
regions
hypo
hyper
J
liver hypo DMRs
CpG shore (4.3%)
CpG shelf (8.0%)
CpG island (2.3%)
interCGI (85.4%)
promoter (5.2%)
5'UTR (0.8%)
1-5Kb (11.2%)
intergenic (9.3%)
3'UTR (4.1%)
exon (19.1%)
intron (34%)
intron-exon border (16.4%)
liver hyper DMRs
CpG shore (4.5%)
CpG shelf (4.7%)
CpG island (1.5%)
interCGI (89.4%)
5'UTR (1.1%)
promoter (4.3%)
1-5Kb (10.5%)
intergenic (13.2%)
3'UTR (3.7%)
exon (18.1%)
intron (34.2%)
intron-exon border (15%)
K
liver
5mC signal
genome coverage
-3kb
TSS
TES
3kb
N/S
young
old

Figure S3

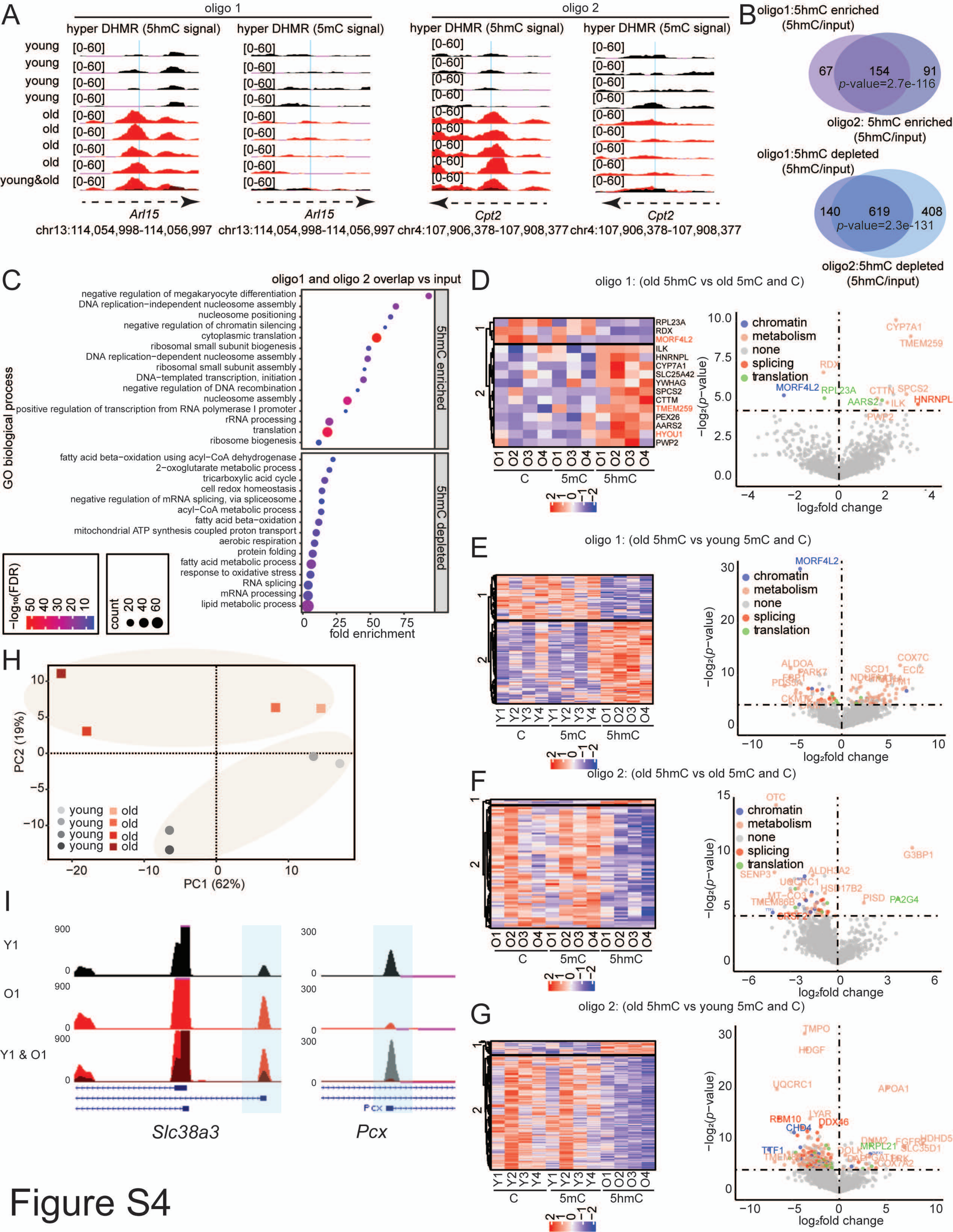
A
oligo 1
oligo 2
hyper DHMR (5hmC signal)
hyper DHMR (5mC signal)
young
old
young&old
Arl15
chr13:114,054,998-114,056,997
Cpt2
chr4:107,906,378-107,908,377
B
oligo1:5hmC enriched (5hmC/input)
67 154 91
p-value=2.7e-116
oligo2: 5hmC enriched (5hmC/input)
oligo1:5hmC depleted (5hmC/input)
140 619 408
p-value=2.3e-131
oligo2:5hmC depleted (5hmC/input)
C
oligo1 and oligo 2 overlap vs input
GO biological process
5hmC enriched
5hmC depleted
fold enrichment
−log10(FDR)
count
D
oligo 1: (old 5hmC vs old 5mC and C)
E
oligo 1: (old 5hmC vs young 5mC and C)
F
oligo 2: (old 5hmC vs old 5mC and C)
G
oligo 2: (old 5hmC vs young 5mC and C)
chromatin
metabolism
none
splicing
translation
log2fold change
−log2(p-value)
H
PC1 (62%)
PC2 (19%)
I
Slc38a3
Pcx
Figure S4

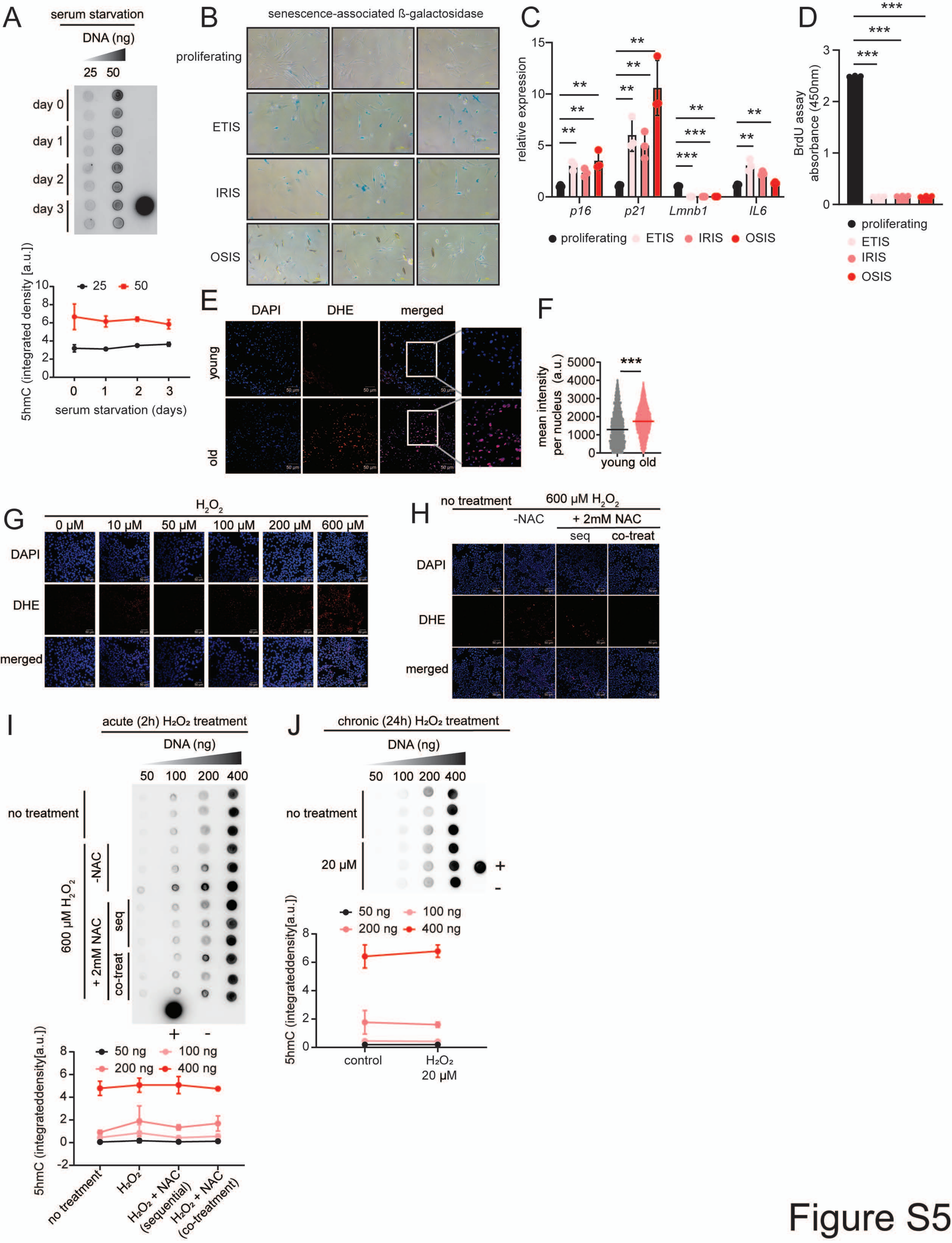

A
serum starvation
DNA (ng)
25 50
day 0
day 1
day 2
day 3
5hmC (integrated density [a.u.])
25
50
serum starvation (days)
B
senescence-associated ß-galactosidase
proliferating
ETIS
IRIS
OSIS
C
relative expression
p16
p21
Lmnb1
IL6
proliferating
ETIS
IRIS
OSIS
D
BrdU assay absorbance (450nm)
proliferating
ETIS
IRIS
OSIS
E
DAPI
DHE
merged
young
old
F
mean intensity per nucleus (a.u.)
young old
G
H2O2
0 µM
10 µM
50 µM
100 µM
200 µM
600 µM
DAPI
DHE
merged
H
no treatment
600 µM H2O2
-NAC
+ 2mM NAC
seq
co-treat
DAPI
DHE
merged
I
acute (2h) H₂O₂ treatment
DNA (ng)
50 100 200 400
no treatment
600 µM H2O2
-NAC
+ 2mM NAC
seq
co-treat
5hmC (integrateddensity[a.u.])
50 ng
100 ng
200 ng
400 ng
no treatment
H₂O₂
H₂O₂ + NAC (sequential)
H₂O₂ + NAC (co-treatment)
J
chronic (24h) H₂O₂ treatment
DNA (ng)
50 100 200 400
no treatment
20 µM
5hmC (integrateddensity[a.u.])
50 ng
100 ng
200 ng
400 ng
control
H₂O₂ 20 µM


Figure S5

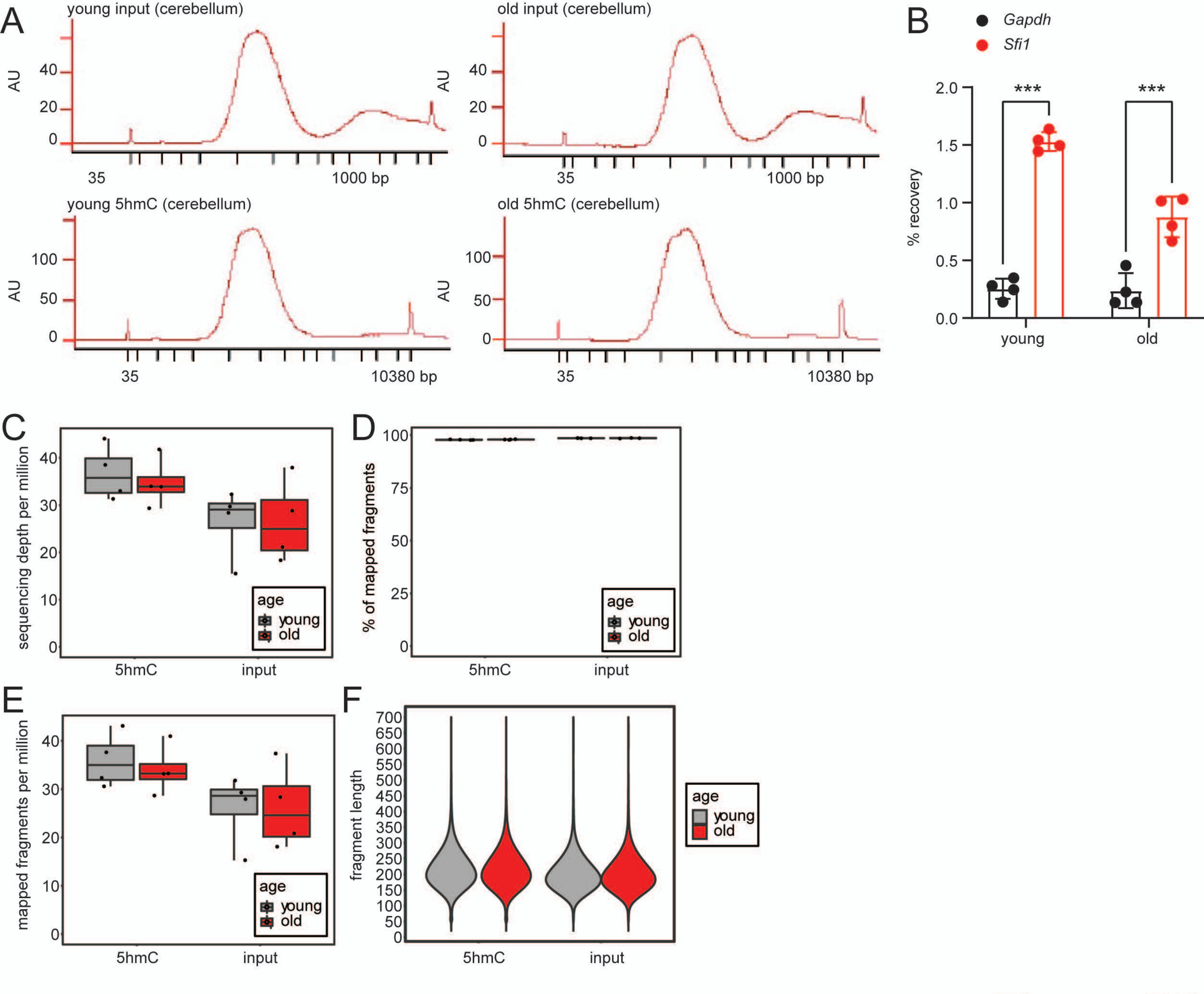
A
young input (cerebellum)
old input (cerebellum)
AU
35
1000 bp
young 5hmC (cerebellum)
old 5hmC (cerebellum)
10380 bp
B
Gapdh
Sfi1
% recovery
***
young
old
C
sequencing depth per million
age
young
old
5hmC
input
D
% of mapped fragments
E
mapped fragments per million
F
fragment length

Figure S6

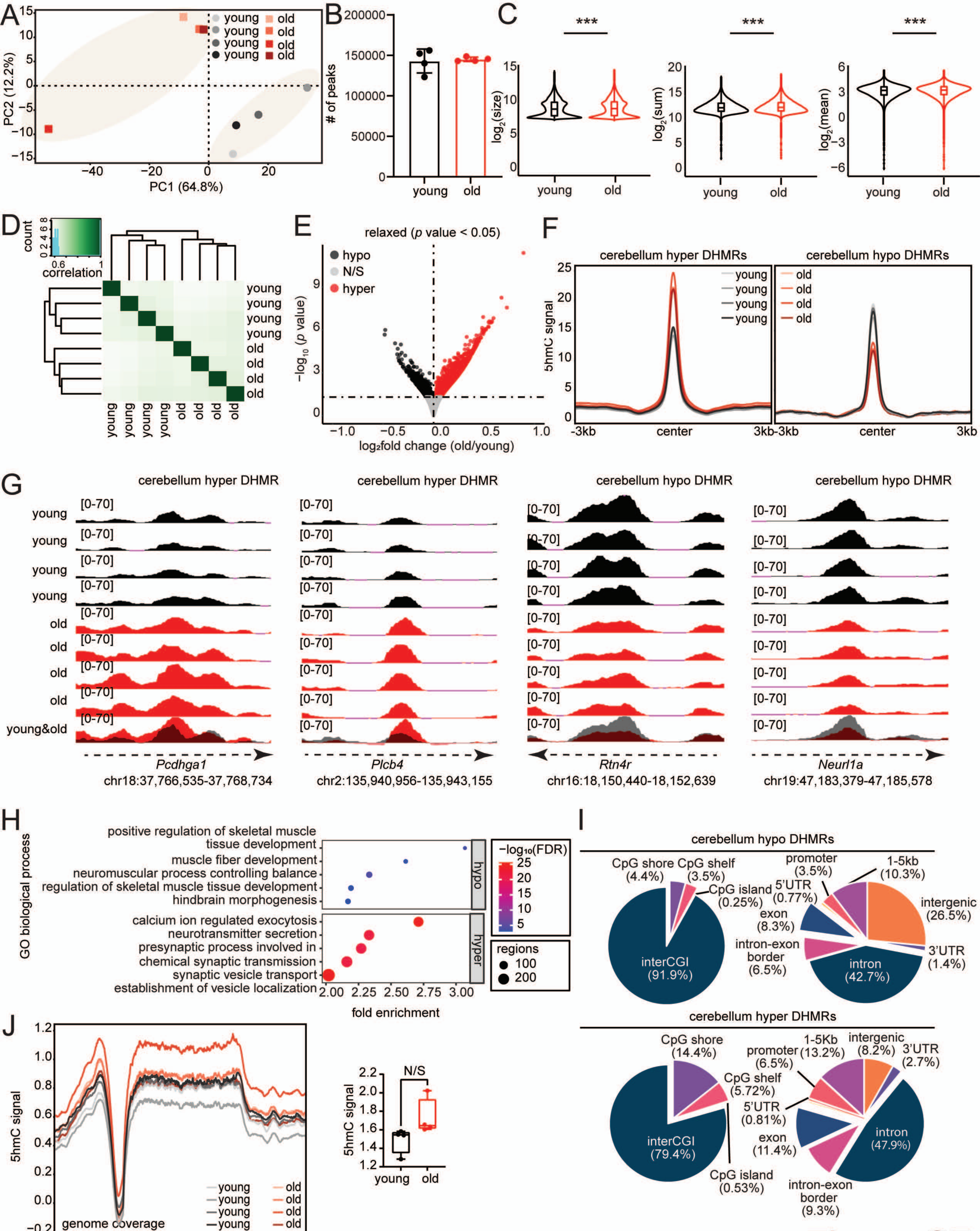
A
B
C
D
E
F
G
H
I
J
PC1 (64.8%)
PC2 (12.2%)
young
old
### of peaks
log2(size)
log2(sum)
log2(mean)
***
count
correlation
relaxed (p value < 0.05)
hypo
N/S
hyper
−log10 (p value)
log2fold change (old/young)
cerebellum hyper DHMRs
cerebellum hypo DHMRs
5hmC signal
-3kb
center
3kb
cerebellum hyper DHMR
cerebellum hypo DHMR
[0-70]
young&old
Pcdhga1
chr18:37,766,535-37,768,734
Plcb4
chr2:135,940,956-135,943,155
Rtn4r
chr16:18,150,440-18,152,639
Neurl1a
chr19:47,183,379-47,185,578
GO biological process
positive regulation of skeletal muscle tissue development
muscle fiber development
neuromuscular process controlling balance
regulation of skeletal muscle tissue development
hindbrain morphogenesis
calcium ion regulated exocytosis
neurotransmitter secretion
presynaptic process involved in chemical synaptic transmission
synaptic vesicle transport
establishment of vesicle localization
fold enrichment
−log10(FDR)
regions
CpG shore (4.4%)
CpG shelf (3.5%)
CpG island (0.25%)
interCGI (91.9%)
promoter (3.5%)
1-5kb (10.3%)
5'UTR (0.77%)
exon (8.3%)
intergenic (26.5%)
intron-exon border (6.5%)
3'UTR (1.4%)
intron (42.7%)
CpG shore (14.4%)
CpG shelf (5.72%)
CpG island (0.53%)
interCGI (79.4%)
promoter (6.5%)
1-5Kb (13.2%)
intergenic (8.2%)
3'UTR (2.7%)
5'UTR (0.81%)
exon (11.4%)
intron (47.9%)
intron-exon border (9.3%)
genome coverage
TSS
TES
N/S

Figure S7

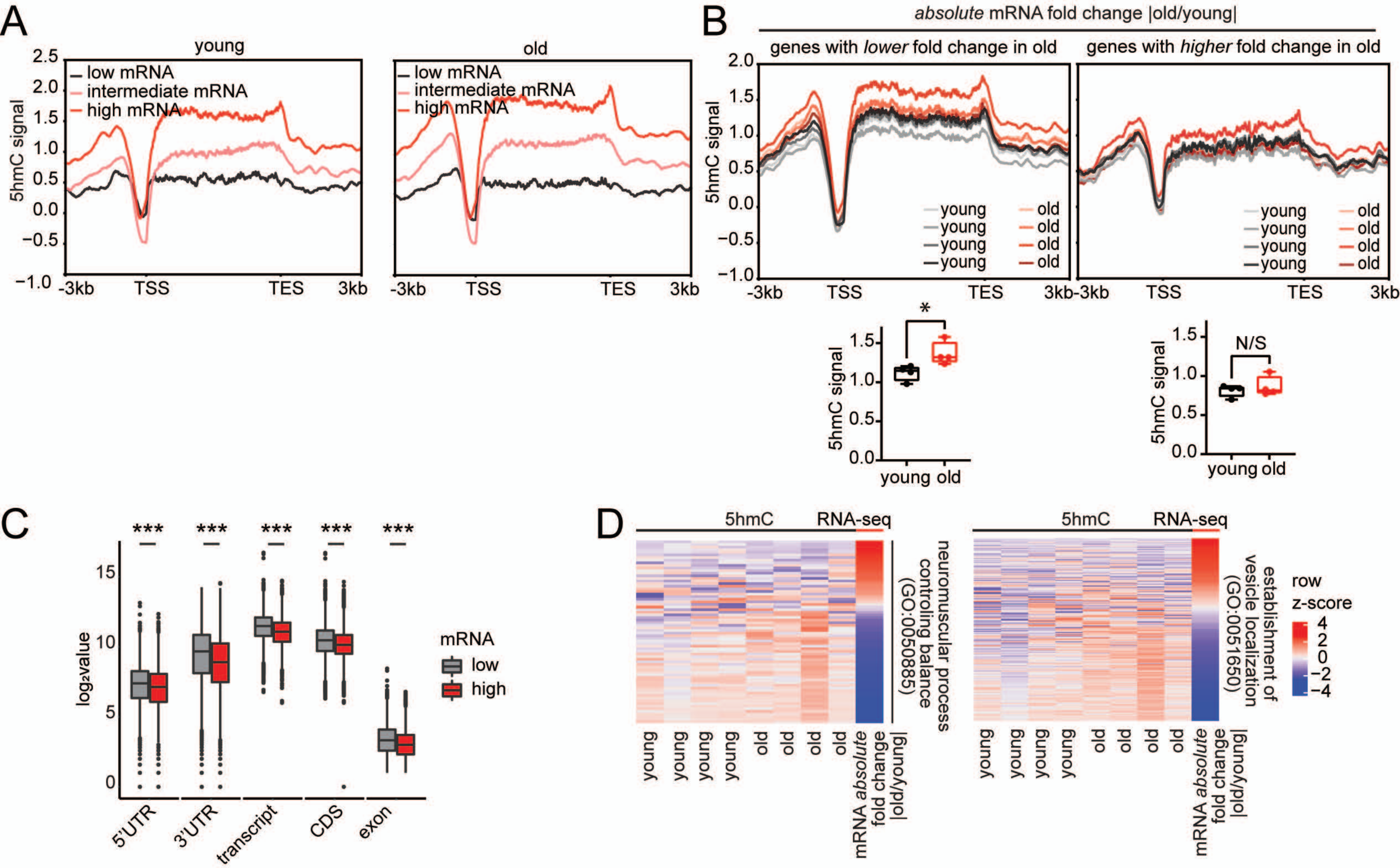
A
young
old
low mRNA
intermediate mRNA
high mRNA
5hmC signal
-3kb
TSS
TES
3kb
B
absolute mRNA fold change |old/young|
genes with lower fold change in old
genes with higher fold change in old
young
old
*
N/S
C
log₂value
mRNA
low
high
***
5'UTR
3'UTR
transcript
CDS
exon
D
5hmC
RNA-seq
neuromuscular process controling balance (GO:0050885)
establishment of vesicle localization (GO:0051650)
mRNA absolute fold change |old/young|
row z-score

Figure S8

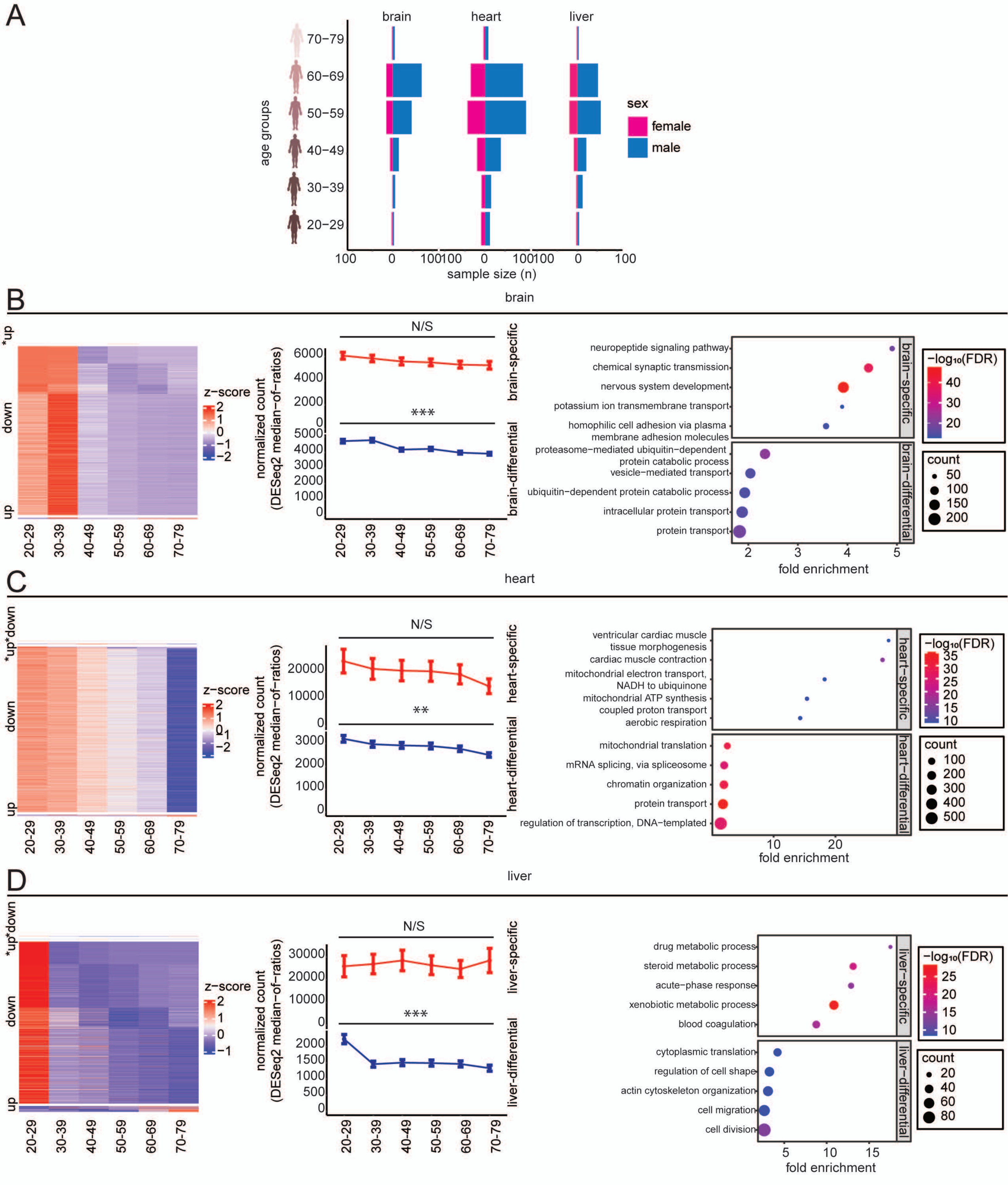
A
brain
heart
liver
age groups
70−79
60−69
50−59
40−49
30−39
20−29
sex
female
male
sample size (n)
B
brain
z−score
normalized count (DESeq2 median−of−ratios)
N/S
***
brain−specific
brain−differential
neuropeptide signaling pathway
chemical synaptic transmission
nervous system development
potassium ion transmembrane transport
homophilic cell adhesion via plasma membrane adhesion molecules
proteasome−mediated ubiquitin−dependent protein catabolic process
vesicle−mediated transport
ubiquitin−dependent protein catabolic process
intracellular protein transport
protein transport
fold enrichment
−log10(FDR)
count
C
heart
N/S
**
heart−specific
heart−differential
ventricular cardiac muscle tissue morphogenesis
cardiac muscle contraction
mitochondrial electron transport, NADH to ubiquinone
mitochondrial ATP synthesis coupled proton transport
aerobic respiration
mitochondrial translation
mRNA splicing, via spliceosome
chromatin organization
protein transport
regulation of transcription, DNA−templated
D
liver
N/S
***
liver−specific
liver−differential
drug metabolic process
steroid metabolic process
acute−phase response
xenobiotic metabolic process
blood coagulation
cytoplasmic translation
regulation of cell shape
actin cytoskeleton organization
cell migration
cell division

Figure S9

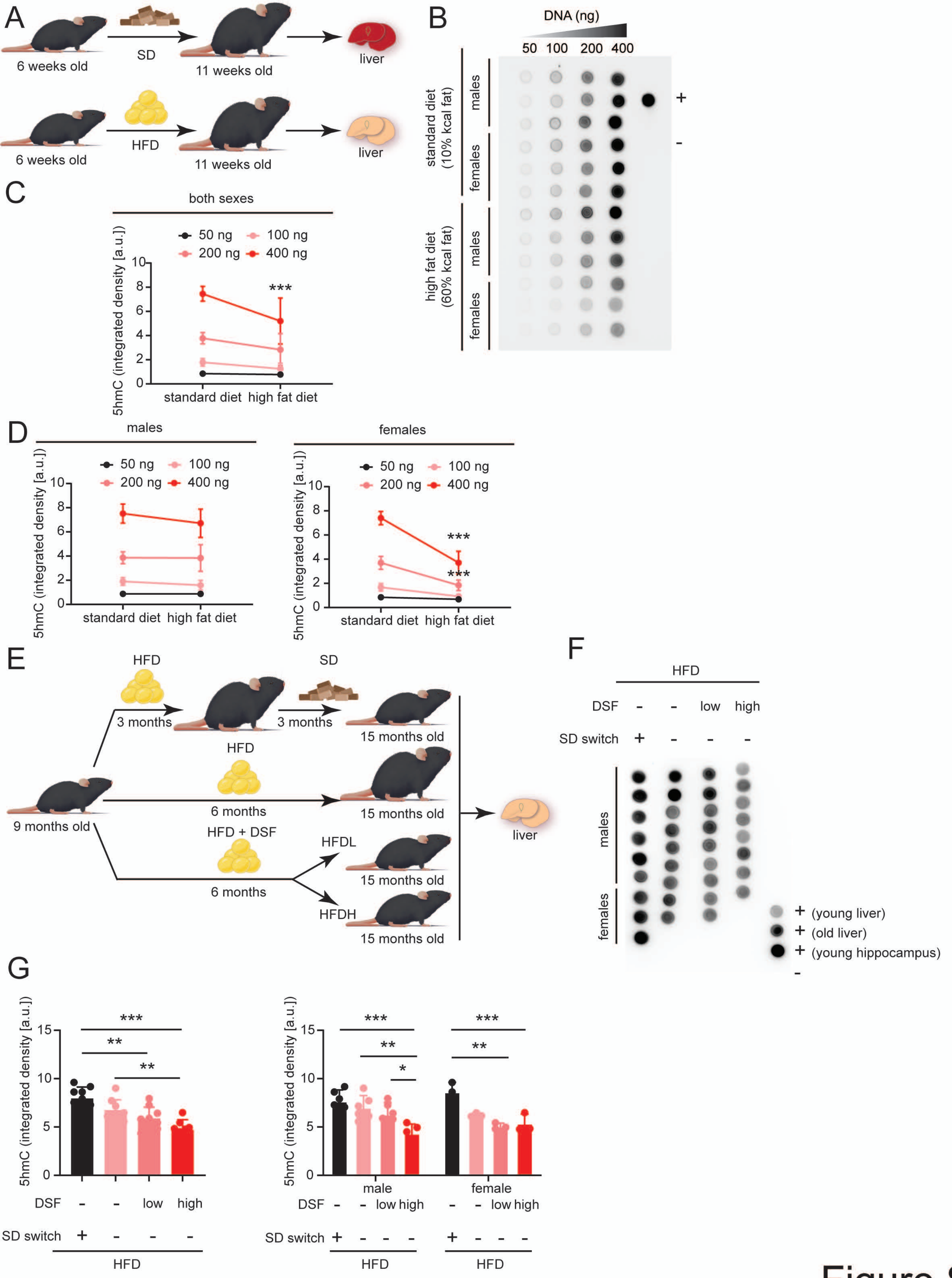

A
6 weeks old
SD
11 weeks old
liver
6 weeks old
HFD
11 weeks old
liver
B
DNA (ng)
50 100 200 400
standard diet (10% kcal fat)
high fat diet (60% kcal fat)
males
females
males
females
+
-
C
both sexes
50 ng 100 ng 200 ng 400 ng
5hmC (integrated density [a.u.])
***
standard diet high fat diet
D
males
females
50 ng 100 ng 200 ng 400 ng
5hmC (integrated density [a.u.])
***
***
standard diet high fat diet
E
HFD
SD
3 months
3 months
15 months old
HFD
6 months
15 months old
9 months old
HFD + DSF
HFDL
15 months old
6 months
HFDH
15 months old
liver
F
HFD
DSF - - low high
SD switch + - - -
males
females
+ (young liver)
+ (old liver)
+ (young hippocampus)
-
G
5hmC (integrated density [a.u.])
***
**
**
*
DSF - - low high
SD switch + - - -
HFD
male
female
DSF - - low high - - low high
SD switch + - - - + - - -
HFD HFD


Figure S10
